# Quasi-species evolution maximizes genotypic reproductive value (not fitness or flatness)

**DOI:** 10.1101/2021.02.23.432496

**Authors:** Matteo Smerlak

## Abstract

Growing efforts to measure fitness landscapes in molecular and microbial systems are motivated by a longstanding goal to predict future evolutionary trajectories. Sometimes under-appreciated, however, is that the fitness landscape and its topography do not by themselves determine the direction of evolution: under sufficiently high mutation rates, populations can climb the closest fitness peak (survival of the fittest), settle in lower regions with higher mutational robustness (survival of the flattest), or even fail to adapt altogether (error catastrophes). I show that another measure of reproductive success, Fisher’s *reproductive value*, resolves the trade-off between fitness and robustness in the quasi-species regime of evolution: to forecast the motion of a population in genotype space, one should look for peaks in the (mutation-rate dependent) landscape of genotypic reproductive values—whether or not these peaks correspond to local fitness maxima or flat fitness plateaus. This new landscape picture turns quasi-species dynamics into an instance of non-equilibrium dynamics, in the physical sense of Markovian processes, potential landscapes, entropy production, etc.

## Introduction

Darwinian micro-evolution is often conceptualized as the motion of populations in the space of all possible heritable types graded by their expected offspring number, the *fitness landscape* [1, 2, 3]. In Wright’s vivid words, the interaction of selection and variation enables populations to “continually find their way from lower to higher peaks” [4], in an open-ended exploration of possible forms that is the hallmark of biological evolution [5]. Thanks to modern sequencing technologies, fitness landscapes have now been measured in a variety of real molecular [6], viral [7] or microbial [8] systems. Thanks to these developments, the goal of *predicting* evolution no longer appears wholly out of reach [9, 10, 8, 11, 12]. In essence, if we know the topography of the fitness landscape—the location of its peaks, valleys, ridges, etc.—we should be able to compute how a population is likely to evolve next. Making such predictions from high-resolution fitness assays, possibly combined with inference methods [13, 14], is a central challenge of quantitative evolutionary theory.

In keeping with Wright’s metaphor, the quantitative analysis of empirical fitness landscapes [15, 16] has so far focused on locating fitness maxima and adaptive walks reaching them in the landscape [17, 18, 13]. This is legitimate in the strong selection regime of evolution [19], where populations are genetically homogeneous except for rare, fitness-increasing fixation events. But in large population subject to high mutational loads, e.g. in RNA viruses, these assumptions usually do not hold, and evolutionary dynamics is better studied within the framework of quasi-species theory [20], where new effects arise.

A hallmark of quasi-species evolution is that populations with different mutation rates experience the same fitness landscape differently. As is well known, populations whose mutation rate are too large can fail to adapt altogether, sometimes (though not always) following sharp “error threshold” phenomena [21, 22]. More subtly, mutational robustness tends to increase during neutral quasi-species evolution [23], and can sometimes outweigh reproductive rate as a determinant of evolutionary success (“survival of the flattest”) [24, 25]. In short, quasi-species evolution favours not just fitness, but some trade-off between fitness and robustness.

These phenomena raise fundamental questions regarding the *dynamical* analysis of fitness landscapes: When is flatter better than fitter? Where are the evolutionary attractors in a given landscape with multiple peaks and/or neutrality? What quantity do evolving populations optimize? Can we estimate the time scale before another attractor is visited? More simply, can we predict the future trajectory of an evolving population—say an influenza strain, or an intra-host HIV population—from its current location, the topography of its landscape, and the mutation rate?

**Figure 1:**
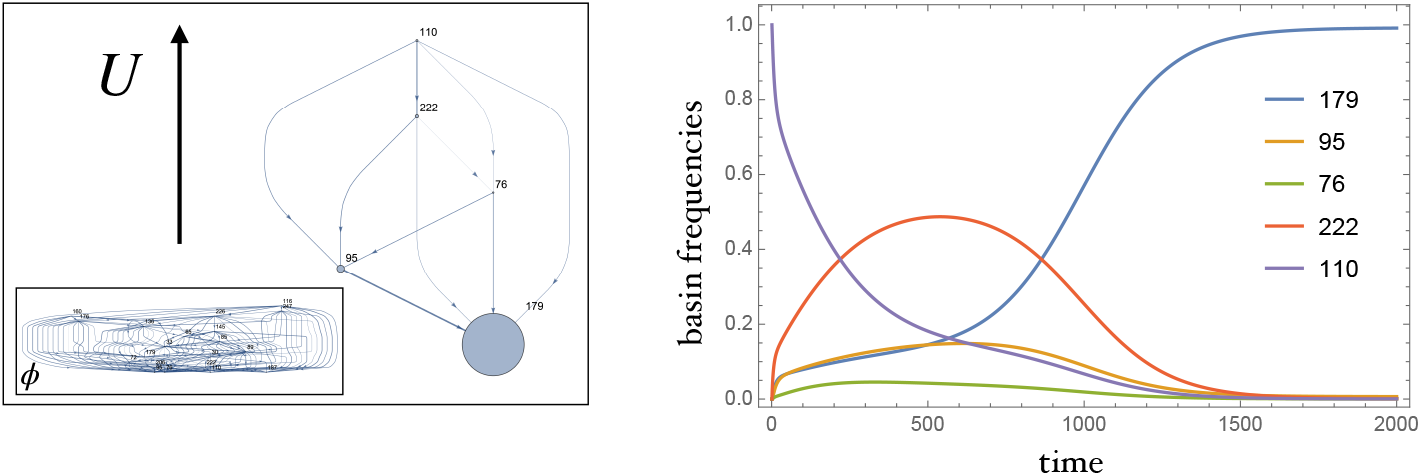
Left: An *NKp* fitness landscape with 20 fitness peaks (inset) is more easily analyzed in terms of local maxima of the reproductive value **R**, which are fewer than fitness peaks, with larger basins of attractions (size of points); see [26] for another illustration of this point. Here the mutation rate is *μ* = 0.1, below the landscape’s error threshold. Right: Solving the Crow-Kimura equation in this landscape shows that a monomorphic population with genotype 110 evolves towards the peak 179 through the basins of attraction of peaks 222 and 95, as dictated by the basin hopping graph of the **R** landscape. Trying to forecast this evolution from the bare fitness landscape **r** itself is hopeless in this case, especially since mean fitness *decreases* over time.

In Ref. [26] I introduced a general framework to coarse-grain and forecast intermittent dynamics in random media. This Letter aims to highlight its applicability to quasi-species evolution, and to explain its connection with (*i*) the concept of *reproductive value* introduced by R. Fisher in relation to age-structured populations [27], and (*ii*) the *retrospective process* discussed in Ref. [28] in the context of mutation-selection balance. Out of these connections emerges a new landscape picture of microevolution, in which large evolving populations climb peaks in the landscape of genotypic reproductive values—not fitness peaks.

## Theory

Quasi-species theory [20] describes the evolution of large molecular or viral populations using ordinary differential equations. To each genotype *i* we associate a density *x*_*i*_(*t*) that changes over time through selection and mutation. In the Crow-Kimura formulation [29], we have a non-linear system of first-order equations

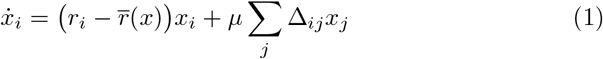

where Δ is the Laplacian matrix of the mutation graph (Δ_*ij*_ = 1 if *i* and *j* can be reached through a mutation, Δ_*ii*_ is minus the number of mutants of *i*, and Δ_*ij*_ = 0 else), *μ* the mutation rate (assumed symmetric for simplicity), *r*_*i*_ is the Malthusian fitness of *i*, and 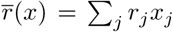 is the population mean fitness. Other formulations are possible, e.g. Eigen’s model where mutation arise as replication events [21]. The key point is that populations are described as a heterogeneous mutant cloud rather than a single consensus sequence. Such a description is necessary for RNA viruses such as influenza or HIV whose mutation rates are so high that any population consists of highly diverse genomes [30].

The quasi-species equation (1) can be linearized by dropping the non-linear mean-fitness term 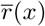, whose sole effect is to maintain normalization of densities. In this way (1) becomes similar to the parabolic Anderson model introduced by Zel’dovich *et al.* as a model of intermittent dynamics [31]; indeed, one salient feature of quasi-species dynamics is its punctuated nature, with populations localizing in regions of the fitness landscapes for long transients before suddenly jumping to remote areas in bursts of rapid adaptation [32].

In [26] I argue that population-dynamical equations of the form 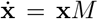 (with **x** = (*x*_*i*_)_*i*_ and *M* = *μ*Δ + *r* in the Crow-Kimura model^1^) are best studied in terms of a related Markov process, generated by the Markov infinitesimal generator

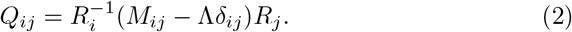

Here **R** = (*R*_*i*_) is the right eigenvector of *M* corresponding to the eigenvalue Λ of *M* with largest real part, normalized so that Σ_*i*_ *R*_*i*_ = 1; in particular **R** depends on the mutation graph and rate. The process (2), which determines **x** as much as the original equation 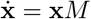, may be interpreted as follows.

Firstly, the component *R*_*i*_ of the right dominant eigenvector measures the contribution of a type *i* to the asymptotic population **x**(*t*) with *t* → ∞: after a long time, the number of descendants of an ancestor of type *i* is proportional to *R*_*i*_. In this sense, *R*_*i*_ is a generalization of Fisher’s *reproductive value*, a notion he originally introduced in age-structured populations as the answer to the question “To what extent will persons of this age, on the average, contribute to the ancestry of future generations?” [27] In the context of quasi-species dynamics, *i* is a genotype and *R*_*i*_ measures the relative number of *i*-clones in a lineage started from an *i*-ancestor. (This should not be confused with the *quasi-species distributions*, given by the left dominant eigenvector **L** = (*L*_*i*_) of *M* and describing the composition of the population at mutation-selection equilibrium; **L** and **R** only coincide if *M* is symmetric.)

Secondly, the process (2) can be interpreted as a *retrospective process* generating the transitions between types observed along successful lineages of a branching process with mean matrix *M*. This process is built as follows: sample an individual at some late time from the equilibrium quasi-species distribution **L**, and trace its ancestry back in time along its genealogical tree. Along this backward path ancestors have genotypes *i*_1_, *i*_2_, …. The retrospective process generates these transitions forward in time [33, 34].^2^.

Thirdly, because (2) generates a Markov process, we can study it using the same tools as other non-equilibrium systems in physics or biology. In particular, we can picture (2) as describing a random walk (with unit diffusivity) in the potential *U*_*i*_ = −2 log *R*_*i*_, characterize the distance to equilibrium using the relative entropy with respect to the Gibbs distribution *π*_*i*_ = *R*_*i*_*L*_*i*_ (the equilibrium distribution for (2)), or measure the irreversibility of evolutionary dynamics with path entropies *s*(*ω*) = log Prob(*ω*)/Prob(*ω*′), where *ω* = (*i*_0_, *i*_1_, …) is a path of the retrospective process and *ω*′ the time-reversed trajectory.

Perhaps more importantly, this new landscape picture provides a solution to the prediction problem in quasi-species dynamics: to find the region of genotype space where a population will go from its current location *i*, find the closest **R** peak *α*. Once the population enters the basin of attraction of that peak, it will concentrate there for a long time—until it jumps to another peak surround by another basin of attraction. In other words, coarse-grained evolutionary trajectories can be defined by aggregating the densities within the basins of attraction of reproductive value maxima, and transitions (punctuations) forecasted in terms of neighboring basins and lowest saddles between them.

## Illustration

To illustrate the predictive power of this principle in a model fitness landscape exhibiting both ruggedness and neutrality, I considered an instance of the *NKp* landscape [35], in which the fitness of a binary sequence of length *N* derives from *N* additive contributions whose values depend on the alleles at *K* linked loci, with a probability *p* that each contribution is exactly zero.^3^. The landscape in Fig. 1, chosen with *N* = 8, epistasis parameter *K* = 6 and neutrality parameter *p* = 0.7, has 20 local maxima and an error threshold at *μ*_*c*_ ≃ 0.2. Comparing the basin hopping graphs of the fitness landscape **r** and of the genotypic reproductive value **R** reveals that most of the complexity of the former is spurious. Moreover, coarse-grained evolutionary trajectories, described by the basin frequencies, are consistent with the succession of transitions predicted from the basin hopping graph [36] of **U**: a population initially concentrated around the genotype 110 (a global fitness maximum) will evolve towards the flatter genotype 179 via the basins of 222 and 95. This simple example highlights the usefulness of studying quasi-species dynamics in terms of reproductive values: while forecasting the coarse-grained dynamics through **R** is straightforward, the same task becomes exceedingly impractical from the peak, valley and plateau structure of the fitness landscape **r**.

## Discussion

A widely shared understanding of the role of mutations in evolution has them feeding raw material to the fitness-maximizing sieve of natural selection. But when mutation rates are high, as they are in *e.g.* RNA viruses [37] and likely were in early life [20], evolutionary success requires more than the discovery of a high-fitness mutant genotype: the mutants of the new mutant must also have relatively high fitness, *i.e.* the mutant type must be mutationally robust. Fisher’s notion of reproductive value combines fitness and flatness into a derived landscape that determines evolutionary trajectories across the spectrum of mutation rates.

The importance of reproductive values in evolutionary dynamics—in particular, its role as the target of selection—has been emphasized repeatedly since Fisher, see e.g. [38, 39, 40]. Usually, though, reproductive values are defined with respect to classes other than genotype: sex, age, etc. The conclusion of this work is that reproductive values defined with respect to genotypes themselves (and parametrized by mutation rates) help us coarse-grain, predict and generally study quasi-species dynamics (*e.g.* using physics-inspired methods) in ways that the bare fitness landscape cannot.

Funding for this work was provided by the Alexander von Humboldt Foundation in the framework of the Sofja Kovalevskaja Award endowed by the German Federal Ministry of Education and Research.

## Footnotes

1 In general *M* is an irreducible, essentially non-negative matrix.

2 Refs. [33, 34] used the process (2) to characterize mutation-selection equilibria; here, by constrast, I use it to describe the dynamics of a quasi-species away from equilibrium.

3 Specifically, the *NKp* model defines the fitness of a length-*N* binary string 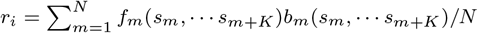 where the fitness components *f*_*m*_ and *bm* are independent with *U* (0, 1) and Bernoulli(*p*) distributions, respectively.

